# P-Selectin promotes SARS-CoV-2 interactions with platelets and the endothelium

**DOI:** 10.1101/2023.02.13.528235

**Authors:** Cesar L. Moreno, Fernanda V. S. Castanheira, Alberto Ospina Stella, Felicity Chung, Anupriya Aggarwal, Alexander J. Cole, Lipin Loo, Alexander Dupuy, Yvonne Kong, Lejla Hagimola, Jemma Fenwick, Paul Coleman, Michelle Willson, Maxwell Bui-Marinos, Daniel Hesselson, Jennifer Gamble, Freda Passam, Stuart Turville, Paul Kubes, G. Gregory Neely

## Abstract

COVID-19 causes a clinical spectrum of acute and chronic illness and host / virus interactions are not completely understood^1,2^. To identify host factors that can influence SARS-CoV-2 infection, we screened the human genome for genes that, when upregulated, alter the outcome of authentic SARS-CoV-2 infection. From this, we identify 34 new genes that can alter the course of infection, including the innate immune receptor P-selectin, which we show is a novel SARS-CoV-2 spike receptor. At the cellular level expression of P-selectin does not confer tropism for SARS-CoV-2, instead it acts to suppress infection. More broadly, P-selectin can also promote binding to SARS-CoV-2 variants, SARS-CoV-1 and MERS, acting as a general spike receptor for highly pathogenic coronaviruses. P-selectin is expressed on platelets and endothelium^3^, and we confirm SARS-CoV-2 spike interactions with these cells are P-selectin-dependent and can occur under shear flow conditions. *In vivo*, authentic SARS-CoV-2 uses P-selectin to home to airway capillary beds where the virus interacts with the endothelium and platelets, and blocking this interaction can clear vascular-associated SARS-CoV-2 from the lung. Together we show for the first time that coronaviruses can use the leukocyte recruitment system to control tissue localization, and this fundamental insight may help us understand and control highly pathogenic coronavirus disease progression.

## Main

The emergence of SARS-CoV-2, the virus responsible for COVID-19, has presented a significant global health challenge. Despite the development of vaccines, the pandemic continues to spread at high frequency globally. Whilst vaccination has significantly reduced disease severity and deaths, the continued high circulation globally has led to in excess of 1 million deaths over the last 12 months. Moreover, at the time of this writing the restriction lifting in China is estimated to result in an additional million deaths this year^4,5^. SARS-CoV-2 infection can cause a range of symptoms, from mild to severe. The main cause of death from COVID-19 is respiratory failure^1^, but other common complications include multi-organ failure^6^, as well as thrombotic events^7^. To understand COVID-19 disease, and to develop more effective therapies, we require a fundamental molecular understanding of how SARS-CoV-2 interacts with the host.

### Host factors sufficient for resistance to authentic SARS-CoV-2

To identify human genes that can modify SARS-CoV-2 infection, we used a whole genome CRISPR activation (CRISPR-SunTag system) strategy (**Fig. 1a**). Cells were transduced with a CRISPRa activation sgRNA lentivirus library and then infected with an early clade SARS-CoV-2 clinical isolate at a lethal dose of 90%. Surviving cells were lysed and integrated sgRNA sequences amplified by PCR and sequenced. The ratio of each sgRNA in control vs. SARS-CoV-2-infected cells was determined and overrepresented guides, i.e. guides that may confer resistance to SARS-CoV-2, were identified using a mixed hierarchical statistical model optimised for whole-genome CRISPR-activation/inhibition screens^12^. In this analysis, guide distribution is evaluated compaired to negative control sgRNA guide distribution, where deviation from control distribution is indicative of functional selection. Our quality control distributions based on negative sgRNA counts showed normally distributed curves (**Supplementary Fig. 1b and c**) and we identified multiple enriched sgRNAs, with the majority of these hits (99.7%) showing a local False Discovery Rate (locFDR) >0.4 (**Fig. 1b and c**, and **Supplementary Table 1a**). Conversely, we did not detect statistically significant sgRNA promoting sensitivity to live SARS-CoV-2 (**Supplementary Table 1b**). Bioinformatic Analysis of the top 100 sgRNA-targeted genes identified significant enrichments in canonical pathways involved in atherosclerosis, ubiquitination, and influenza infection **(Fig. 1d)**. Additional characterization of our top resistance genes using KEGG highlighted that the main molecular processes leading to protection involves protein glycosylation (**Fig. 1e**).

**Fig 1.**
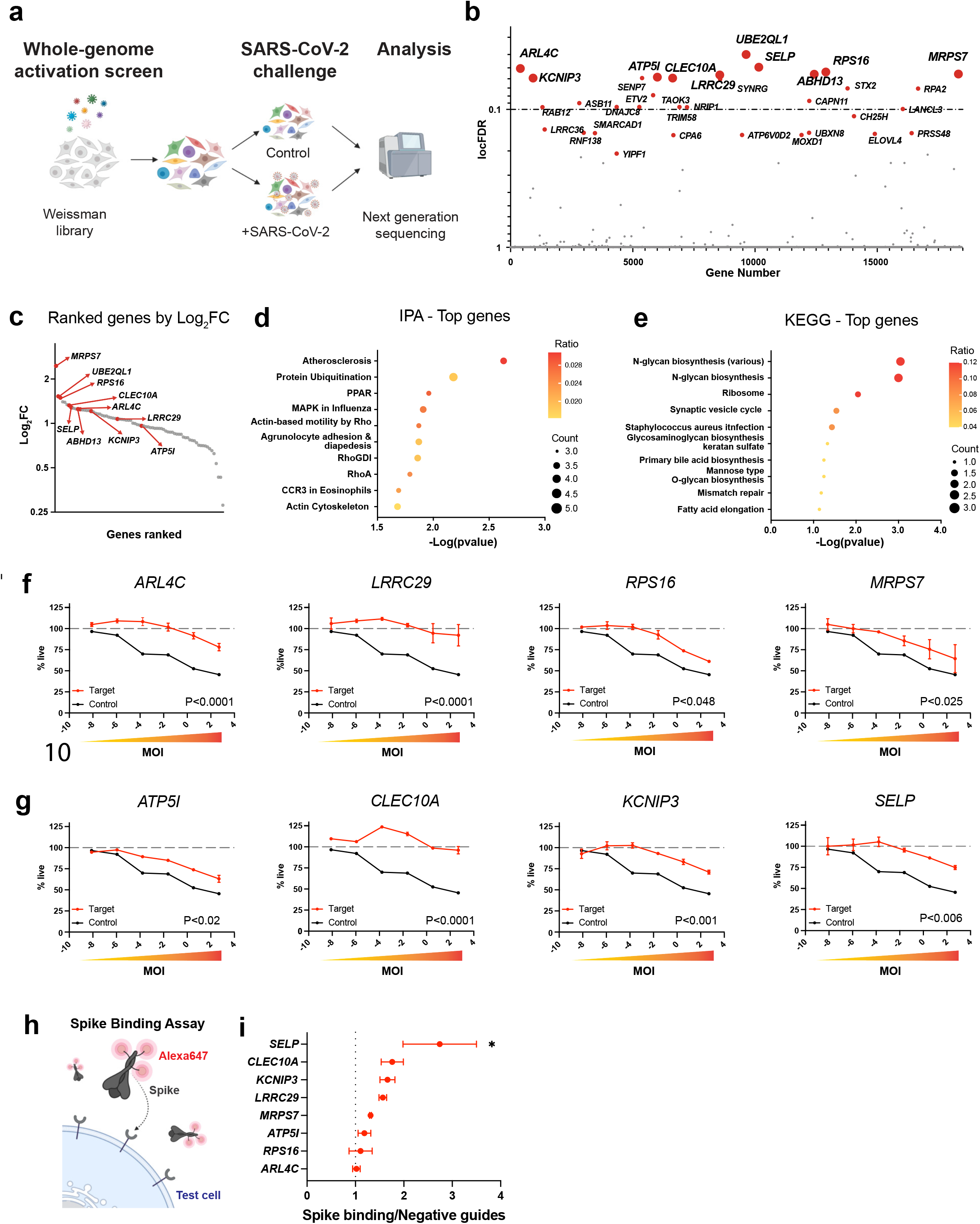
CRISPR-Activation screen against authentic SARS-CoV-2. **a**, Schematic showing Whole-Genome CRISPR-Activation screen in ACE2-HEK293 cells using the Weissman library. Cells were transduced (0.3 MOI) with lentiviral pools encoding individual activation sgRNAs tiling the genome. Cells were then inoculated with authenticSARS-CoV-2 or parallel controls and guides promoting SARS-CoV-2 were identified by sequencing. **b**, Top genes by local FDR (<0.1 considered significant) identified after selection. See also Supplementary Table 1. **c**, Plot showing sgRNA enrichment (log fold change) vs. gene ranking. Top pathways identified using **d**, Ingenuity Pathway Analysis (IPA) or **e**, KEGG pathways. Independent validation of top hits including **f**, membrane proteins **g**, cytosolic (*RPS16*) and mitochondrial proteins (*ATP5I, MRPS7*) promoting SARS-CoV-2 resistance. Significance determined by Simple Linear Regression. **h**, diagram of SARS-CoV-2 FACS spike binding assay. **i**, CRISPRa-driven expression of SELP/ (P-Selectin) promotes binding to SARS-CoV-2 spike protein. Significance obtained by One-way ANOVA and Sydak test, *, P<0.05.

We next independently validated the top SARS-CoV-2 resistance genes using two sgRNAs per candidate. Validated resistance genes were cytosolic proteins, ribosomal proteins, mitochondrial proteins, or cell surface proteins (**Fig. 1f** and **g**). For example, we identified *ARL4C*, encoding a member of the ARF family of GTPases, which participates in various cellular processes including vesicle trafficking and cholesterol export^8^; *LRRC29* (FBXL9P), which is a component of the SKP1-cullin-F-box complex that regulates phosphorylation-dependent ubiquitination^9^; *RPS16*, which encodes a component of the 40s cytoplasmic ribosome^10^; and *MRPS7* which encodes a mitochondrial ribosomal 28S subunit protein^11^. Of the top ranked SARS-CoV-2 resistance genes encoding membrane localized proteins we found *ATP5I*, which encodes ATP Synthase Membrane Subunit E, a component of the mitochondrial f1f0-atp synthase^8^, with plasma membrane localised f1f0 plays a critical role in influenza budding^12^; *CLEC10A*, coding for a type II transmembrane lectin that is involved in immune response and inflammation, and which has been reported to interact with spike protein^13^; *KCNIP3*, a potassium channel interacting protein^14^ that can also act as a Ca^2+^ regulated transcriptional repressor^15^; and *SELP*, which encodes for the protein P-selectin, a critical cell adhesion molecule that also controls fibrin deposition^3,16,17^. We tested if any of these resistance genes were sufficient to promote SARS-CoV-2 spike binding (**Fig. 1h**), and found P-selectin could directly interact with SARS-CoV-2 spike (**Fig. 1i**).

### P-selectin interacts with pathogenic coronavirus spike proteins to protect target cells

P-selectin is expressed on activated platelets and endothelium, where it controls platelet aggregation and leukocyte recruitment to sites of inflammation, especially sites of vascular injury^18,19^. Since inflammation, platelet aggregation, and vascular injury are all hallmarks of COVID-19, we considered the role of P-selectin in SARS-CoV-2 biology. To confirm our CRISPRa results, we used cDNA to express *GFP* control, *ACE2*, or *SELP* and then evaluate impact on SARS-CoV-2 (**Fig. 2a**). Indeed, ectopic expression of SELP (or ACE2) was sufficient to confer SARS-CoV-2 spike binding (**Fig. 2b, Supplementary Fig. 2a**, quantified in **Fig. 2c**). We next assessed if SELP/SARS-CoV-2 spike interactions were sufficient to allow SARS-CoV-2 infection. To this end, we generated a DsRED-SARS-CoV-2-Pseudovirus expressing truncated spike protein^20^. While pseudovirus could infect *ACE2* expressing cells, *SELP* expressing cells did not support pseudovirus infection (**Fig. 2d**, quantified in **e**). Authentic SARS-CoV-2 infects cells primarily via an ACE2/TMPRSS2 entry pathway^21^. Since P-selectin can antagonise both pseudovirus and authentic infection of ACE2 expressing cells, we hypothesised that it would also block infection of ACE2/TMPRSS2+ target cells. Accordingly, compared to ACE2/TMPRSS2 cells transfected with GFP control vector, ACE2/TMPRSS2/SELP+ cells showed a dose-dependent protection from authentic SARS-CoV-2 infection (**Fig. 2f**, D614G, **Fig. 2g**, Delta).

**Fig 2.**
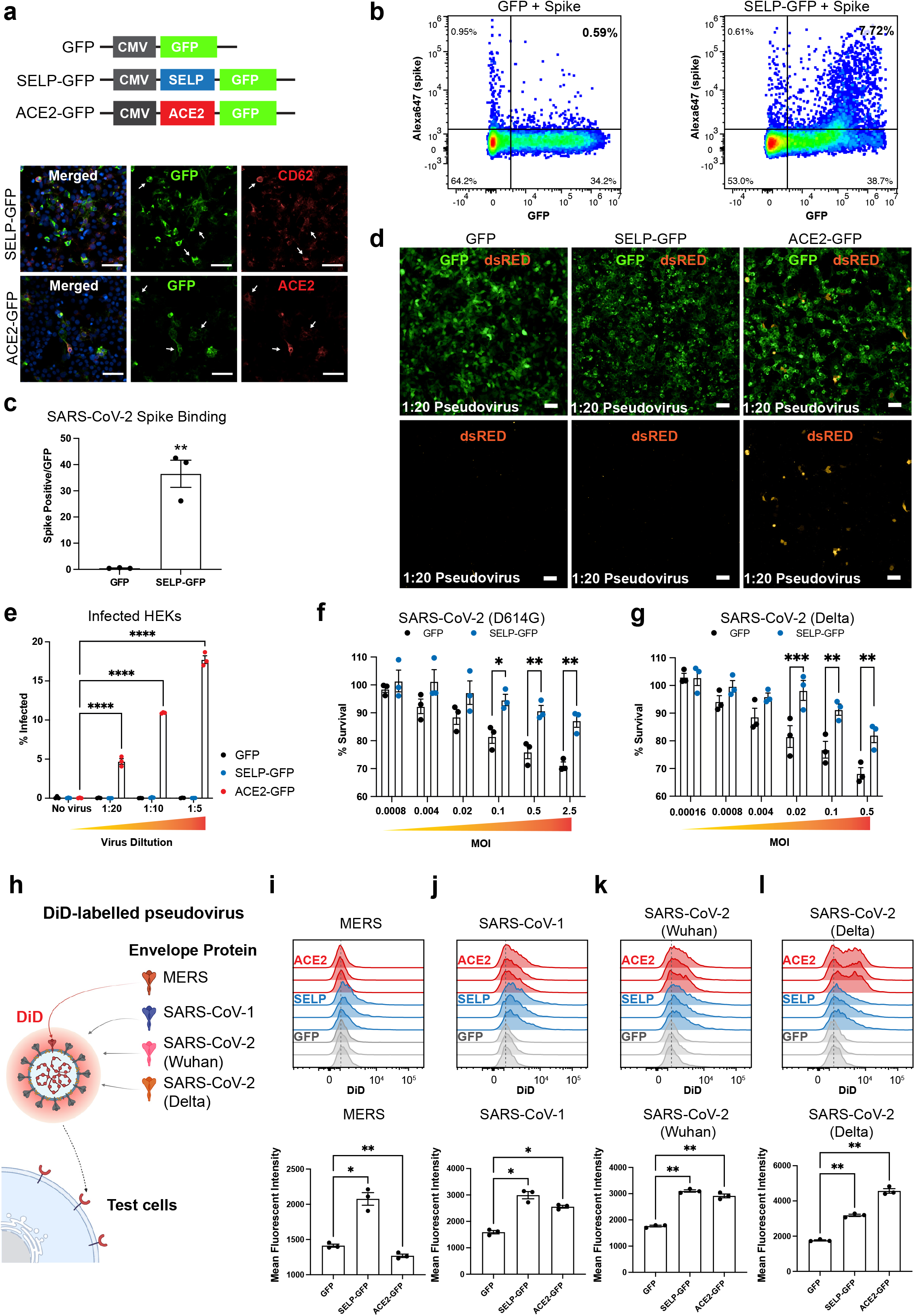
P-Selectin interaction with pathogenic coronavirus spike proteins. **a**, Ectopic ACE2 or SELP expression strategy. **b**, Representative flow-cytometry plots showing binding of P-Selectin to SARS-CoV-2 spike protein. **c**, Quantification of **b**. Significance was determined by paired t-test, **P<0.01. **d**, Images of ACE2 or SELP expressing cells infected with SARS-CoV-2 spike-expressing pseudolentivirus. **e**, Quantification of **d**, significance was determined by 2-way ANOVA and Dunnett test, ****P<0.001. Infection of ACE2/TMPRSS2-expressing cells with authentic SARS-CoV-2. **f**, D614G or **g**, delta variants are suppressed by co-expression of SELP. Significance determined by 2-Way ANOVA and Sydak test. **h**, Schematic of DiD-labelled pseudolentivirus assay used in **i-l. i**, Flow cytometry histograms and **j**, quantification of binding intensity presented for pseudovirus expressing MERS, SARS-CoV-1, SARS-CoV-2 (wuhan), and SARS-CoV-2 (Delta) variants. Significance was determined by 1-way ANOVA and Dunnett test, *P<0.05, **P<0.01.

P-selectin is a lectin that binds to glycoproteins and it is possible that P-selectin shows broad specificity for other gycosylated coronavirus spike proteins. To test this, we generated individual pseudoviruses expressing spike proteins from highly pathogenic members of the coronaviridae family. To visualize interactions, we labelled pseudoviruses with the lipophilic fluorescent probe DiD^4^ and tested if expressing P-selectin could promote cell/pseudovirus interactions (**Fig. 2h**). As reported^22,23^, ACE2 can bind SARS-CoV-1 and SARS-CoV-2 spike, but not MERS spike, whereas P-selectin can promote binding to both SARS and MERS spike proteins. Moreover, as described by others^24^, ACE2 showed enhanced affinity for SARS-CoV-2 delta spike, whereas P-selectin showed comparable binding to all pathogenic spike proteins (**Fig. 2i-l**).

### P-selectin controls SARS-CoV-2 spike binding to platelets and endothelial cells

In platelets, P-selectin is found in preformed intracellular stores where it is translocated to the cell surface upon inflammatory or thrombogenic challenges (**Fig. 3a**). Thrombin-activated human primary platelets express cell surface P-selectin (**Fig. 3b**) and bind SARS-CoV-2 spike protein which is blocked by the P-selectin antagonist Fucoidan (**Fig. 3c**, quantified in **d**). Fucoidan could also block interactions between activated platelets and DiD labelled SARS-CoV-2 pseudovirus (**Fig. 3e**). Moreover, platelets that bind SARS predominately express surface P-selectin (**Fig. 3f**, quantified in **g**). P-selectin is also expressed on the activated endothelium, where it can promote leukocyte trafficking to sites of inflammation^25^. Since SARS-CoV-2 spike can interact with P-selectin and SARS-CoV-2 virus is found in inflamed tissue and at sites of vascular injury where P-selectin is also found^7^, we hypothesised that SARS-CoV-2 may use the host leukocyte trafficking system to control its localization during infection. To this end, we plated GFP or SELP-GFP+ cells and then evaluated the ability of P-Selectin to promote SARS-CoV-2 pseudovirus adhesion under venular flow rates (**Fig. 3h**). Indeed, under these conditions, we observed an accumulation of fluorescent SARS-CoV-2 pseudovirus particles in the capillaries plated with SELP-GFP (**Fig. 3i-k and Supplementary Fig. 2b**) or ACE2 (**Supplementary Fig. 2c**) expressing cells, but not GFP control cells. When we quantified the number of foci per cell area, we found a 7-fold increase in SARS-CoV-2 pseudovirus particle adhesion to cells expressing P-selectin (**Fig. 3k**) and this increase was also confirmed by mean fluorescent intensity (**Supplementary Fig. 2b**). To determine whether SARS-CoV-2 pseudovirus binding is dependent on native P-selectin expression, we targeted *SELP* in primary human umbilical vein endothelial cells (HUVECs) (**Fig. 3l**), observing approximately 20% (guide 1) and 40% (guide 2) reduction of P-selectin expression compared to a control guide by immunocytochemistry (**Fig. 3l**) and flow cytometry (**Fig. 3m, Supplementary Fig. 2d**). Importantly, reducing P-selectin expression in HUVECs was sufficient to decrease SARS-CoV-2 spike protein binding to endothelial cells (**Fig. 3n**).

**Fig 3.**
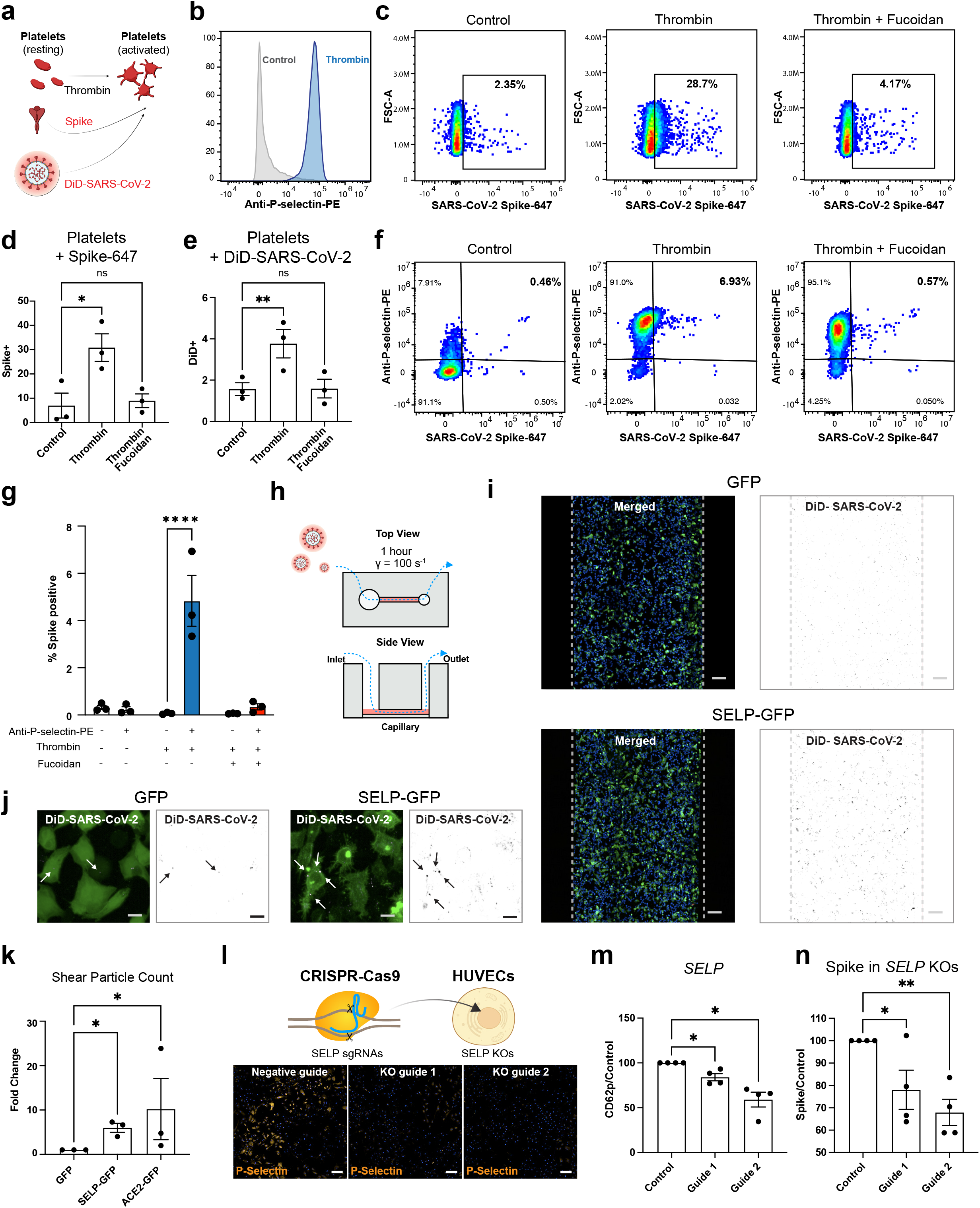
Platelets and endothelium use P-selectin to bind to SARS-CoV-2 spike. **a**, Schematic for testing P-selectin function in platelets. **b**, Thrombin upregulates cell surface P-selectin on platelets. **c**, Thrombin-activated platelets bind SARS-CoV-2 spike protein and this is blocked by the P-selectin antagonist fucoidan. **d**, Quantification of **c**. Significance was determined by 1-way ANOVA and Dunnett test, *P<0.05. **e**, DiD-labelled SARS-CoV-2 pseudovirus bind platelets and this depends on P-selectin. Flow cytometry quantification with significance determined by 1-way ANOVA and Dunnett test, **P<0.05. **f**, Platelets that bind SARS-CoV-2 spike protein are P-selectin positive. **g**, Quantification of **f**, significance was determined by 2-way ANOVA and Sydak test, ****P<0.0001. **h**, Schematic of microfluidic devices used to study SARS-CoV-2 pseudovirus binding to P-selectin under shear conditions. **I**, Representative images of capillary lined with cells expressing GFP or P-selectin-GFP after one hour of flow with SARS-CoV-2 pseudovirus-containing media. Scale bar = 100μm. **j**, Representative images detailing accumulation of labelled pseudovirus particles in GFP or P-selectin-GFP expressing cells. Scale bar = 10μm. **k**, Automated quantification of total DiD area/cells within the capillary. Significance was determined by 1-way ANOVA and Dunnett test, *P<0.05. **l**, CRISPR targeting of SELP in primary endothelial cells (HUVEC) reduces P-selectin staining (Scale bar = 200μm), quantified by flow cytometry in **m**, significance was determined by 1-way ANOVA and Dunnett test, *P<0.05. **n**, CRISPR-targeted SELP-edited HUVEC cells bind less SARS-CoV-2 spike protein, significance was determined by 1-way ANOVA and Dunnett test, *P<0.05, **P<0.01.

### Anti-P-selectin treatment regulates authentic SARS-CoV-2 trafficking *in vivo*

To study the physiological role of P-selectin in SARS-CoV-2 trafficking, we visualised the movement of DiD-labelled SARS-CoV-2 clinical isolate in the infected lung *in vivo*. We confirmed that labelled authentic SARS-CoV-2 viral particles exhibited predicted tropisms for ACE2+ target cells by flow cytometry (**Supplementary Fig. 3a, b**). To monitor the biology of virus trafficking during SARS-CoV-2 infection, we first administered an intranasal dose of the SARS-CoV-2 isolate to transgenic mice expressing human ACE2 (**Fig. 4a**). Four days later, we injected DiD-labelled SARS-CoV-2 i.v. and then imaged the lung vasculature using intravital pulmonary microscopy. We observed that after i.v. injection of DiD-labelled virus, most SARS-CoV-2 particles remained static and bound to (CD31+) endothelium within the vascular capillaries (**Fig. 4b**). However, in some instances moving particles were observed, which displayed intermittent interactions with (CD49b+) platelets (**Fig. 4c**). A plot of individual SARS-CoV-2 particles displays these transitory or stationary modalities (**Supplementary Fig. 3c**). To test the role of P-selectin in these interactions, we injected animals with anti-P-selectin and then evaluated changes in DiD-SARS-CoV-2 localisation. Qualitatively, we observed a rapid mobilisation of DiD-SARS-CoV-2 following anti-P-selectin injection (**Fig. 4d, Supplementary Video 1**). Quantitatively, we measured virus particle intensity at baseline and then tracked these regions over time, observing a rapid and significant loss of viral particles after anti-P-selectin injection (**Fig. 4e)**. An individual quantification of all measured virus particles further supports that anti-P-selectin treatment led to a reduction in vascular interaction **(Fig. 4f)**. The P-selectin-dependent change in signal was computed using one-phase decay and the anti-P-selectin decay constant (K=0.29) was significantly larger than control (K=0.002) (**Fig. 4g**), and based on the rolling average change in intensity, the largest decrease in intensity occurred at the time of treatment administration (**Fig. 4h)**. Further, while control animals exhibit a steady state accumulation of labelled SARS-CoV-2 particles, no new binding of viral particles was detected after animals received blocking P-selectin antibody (**Fig. 4i**). Overall, these data supports that SARS-CoV-2, via its glycosylated spike protein, can interact with P-selectin on both platelets and the endothelium, and through these interactions SARS-CoV-2 can control its physiological localisation within the host.

**Fig 4.**
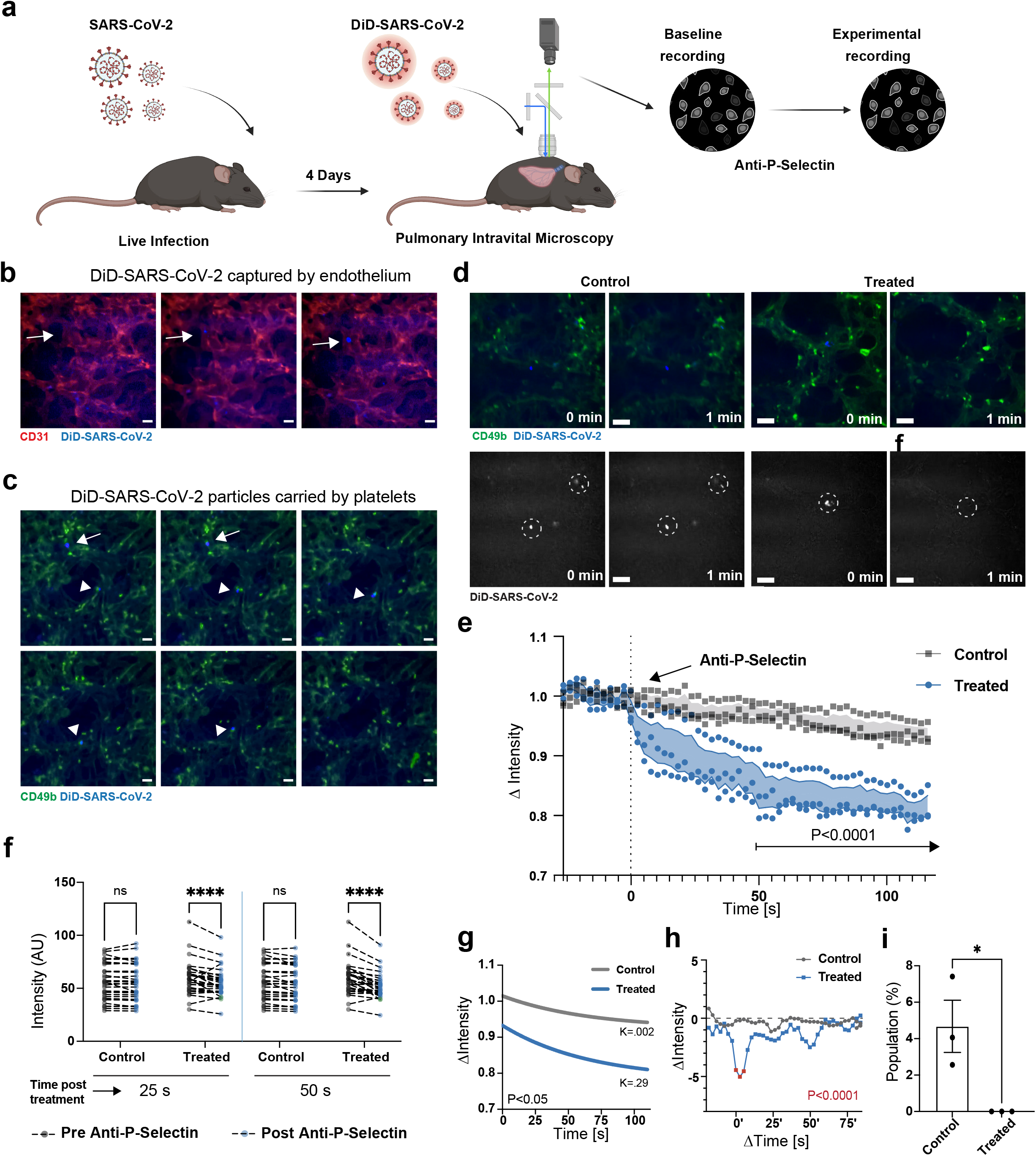
Authentic SARS-CoV-2 uses P-selectin to bind to the endothelium in lung capillary beds. **a**, Schematic of experimental set up. Mice are infected with authentic SARS-CoV-2, then 4 days later, infected mice are injected i.v. with DiD-labelled SARS-CoV-2 and virus trafficking to the lung is monitored by intravital microscopy. Mice are then treated with anti-P-selectin and SAR-CoV-2 interactions with platelets (anti-CD49b-FITC) and the endothelium vasculature are monitored. **b**, Representative time lapse images showing DiD-labelled SARS-CoV-2 binding to the pulmonary endothelium (arrow) (Scale bar = 20 μm). **c**, Representative time lapse images showing travelling (arrow) DiD-labelled SARS-CoV-2 and interacting with platelets (arrowhead) .(Scale bar = 20 μm). **d**, representative time lapse images starting at baseline (Scale bar = 20 μm). DiD-labelled SAR-CoV-2 is blue, platelets (anti-CD49b) are dark green, vasculature is labelled with nonspecific staining (light green). Top panel shows platelet and SARS-CoV-2 signal, bottom panels show only SARS-CoV-2 signal. Circles highlight location of viral particles before or after Anti-P-selectin administration in control and treatment. **e**, quantification of blue (DiD-SARS-CoV-2) fluorescence intensity of the vasculature over time +/- addition of anti-P-selectin antibody. **f**, Intensity measured at regions of interest containing viral particles throughout baseline and post anti-P-selectin introduction. Significance was determined by 2-way ANOVA and Sydak test. Significance was determined by 2-way ANOVA and Sydak test. ****, P<0.0001. **g**, One-phase decay trace showing fluorescent intensity (DiD-SARS-CoV-2) over time decay over time. Significance was determined by sum-of-squares F test. *,P<0.05. **h**, Rate of change over time +/- addition of Anti-P-Selectin antibody. Significance was determined by 2-way ANOVA and Sydak test. **i**, Blinded quantification for the percent of newly bound DiD-SARS-CoV-2 virus after treatment vs. controls. Significance was determined by t test *,P<0.05.

## Discussion

Our team conducted a comprehensive functional analysis of the human genome to identify genes that when upregulated can modify SARS-CoV-2 infection. Through this, we discovered new host factors that can control SARS-CoV-2 infection, information that will help guide our understanding of COVID-19 pathology. We focused on interactions between host anti-SARS-CoV-2 genes and the SARS-CoV-2 spike protein, and found that P-selectin, a cell adhesion molecule expressed on activated platelets and endothelial cells, can bind to pathogenic coronavirus spike proteins and suppress infection. From a physiological perspective, we demonstrated that endogenous P-selectin can also promote binding to SARS-CoV-2 spike, and these interactions can occur under conditions of shear stress. Importantly, *in vivo*, P-selectin in the lung controls the movement of viral particles within the vasculature, providing evidence for the first time that SARS-CoV-2 hijacks our leukocyte recruitment systems to home to inflamed tissues.

From intravital microscopy we observed viral particles interacting with the endothelium, as well as migrating virus particles closely interacting with platelets. These observations are congruent with *ex vivo* reports that SARS-CoV-2 particles accumulate and aggregate in vascular beds^26^ or reports that platelets isolated from COVID-19 patients associate with SARS-CoV-2 RNA^27^. Overall, our research supports the idea that, by facilitating the anchoring of the SARS-CoV-2 virus to the vasculature, and promoting interactions with platelets, P-selectin plays a critical role in SARS-CoV-2 localization during infection. At the cellular level, post-mortem work on COVID-19 patients identified instances of endotheliitis and in some cases SARS-CoV-2 infection in endothelial cells^28^. Although the level of endothelial infection in non-severe COVID-19 remains to be determined, it is possible that endothelial expression of P-selectin could provide some protection from infection or, alternatively, P-selectin could increase virus/vascular interactions and in this way promote endothelial infection. We also provide evidence that surface P-selectin can interact with SARS-CoV-1 and MERS spike proteins, suggesting engagement with the leukocyte trafficking machinery may be a fundamental aspect of coronavirus biology.

The physiological roles of P-selectin in the context of SARS-CoV-2 infection are complex, and likely depend on disease stage, severity, and microenvironment. Multiple studies have investigated associations between P-selectin expression and COVID-19, including correlations with disease severity. Generally most^29–34^, but not all^35–37^, studies show that soluble P-selectin expression increases with disease severity, and serum P-Selectin has shown sensitivity and specificity as a biomarker for Long-COVID^38^. Moreover, a recent trial using the anti-P-Selectin antibody Crizanlizumab to treat 22 COVID-19 patients with moderate disease showed that a single injection of Crizanlizumab was safe^39^ however Crizanlizumab did not alter the clinical outcomes measured (hospital discharge, or clinical status). Of note, this was a small study and neither control nor Crizanlizumab treated patients experienced serious outcomes (ventilation, vascular event, stroke, myocardial infarction), moreover, blocking viral pathways are more effective early in infection, where as in the hospitalisation phase targeting the immune response has been more effective. Building on these efforts a larger trial is currently underway as part of the NIH ACTIV-4a platform. In summary, we report the unexpected finding that the leukocyte adhesion molecule P-selectin can also bind to SARS-CoV-2 spike protein and block infection, and that SARS-CoV-2 can use this interaction to home to sites of inflammation.

## Online content

## Acknowledgements

G.G.N. is supported by grants from the NHMRC, the ARC, and a Philanthropic donation from Dr. John and Anne Chong.. F.V.S.C. is supported by a fellowship from Canadian Institutes of Health Research.

## Author contributions

Screen development and analysis were performed by F. C, A.O.S, A.A, and C.L.M. Target validation using Authentic SARS-CoV-2 was done by A.O.S, A.A, and C.L.M. Spike experiments included the help of L.L and A.C. Platelet and microfluidic experiments were done with the participation of C.L.M, A.D, Y.K, L.H, J.F, and F.P. Endothelial cell experiments were performed by C.L.M and P.C. Animal intravital work was performed by F.V.S.C. with the help of M.W. Vero-E6-hACE2 and Hela cells infection with labelled authentic SARS-CoV-2 was performed by M.B.M. Study analysis was performed by C.L.M, P.K, D.H, S.T, A.O.S, and G.G.N. The manuscript was drafted by C.L.M and G.G.N with all authors providing editorial support. The study was supervised by J.G, F.P, S.T, P.K, and G.G.N,

## Competing Interests

Need competing interest for each author

## Methods

### Lentivirus generation

Relevant vectors were co-transfected with packaging plasmids pCAG-VSVG and psPAX2 Addgene plasmids 35616 and 12260, respectively, using lipofectamine 3000 according to manufacturer’s instructions. Molecular ratios of 3:3:1 corresponding to the vector of interest, psPAX2, and pCAG-VSVG were used to package viruses in HEK293FT cells. These cells were cultured in DMEM (Sigma-Aldrich) medium supplemented with 10% FBS, 1X GlutaMAX, 1X NEAA, and 1X PenStrep. Medium was replaced the day after transfection, and cells were cultured for an additional 24 hours. Medium was collected after this period and filtered through 0.45μm ultra-low protein binding filter (Merck Millipore) to remove cell debris. Medium was then concentrated using PEG (Abcam) according to manufacturer’s instructions. Virus aliquots were stored at -80°C before use.

### Whole genome CRISPR activation sgRNA lentivirus library preparation

The human genome-wide CRISPRa-V2 library (Addgene 1000000091) was co-transfected with packaging plasmids pCAG-VSVG and psPAX2 (Addgene plasmids 35616 and 12260, respectively). Briefly, a T-175 flask of 80% confluent 293LTV cells (Cell Biolabs) was transfected in OptiMEM (Thermo Fisher Scientific) using 8 μg of the plasmid library, 4 μg pCAG-VSVG, 8 μg psPAX2, 2.5 μg pAdVantage (Promega), 30 μl of P3000 Reagent (Thermo Fisher Scientific), and 30 μl of Lipofectamine 3000 (Thermo Fisher Scientific). Cells were incubated overnight and then media was changed to DMEM (Sigma-Aldrich) with 10% FBS and 1x GlutaMAX (Thermo Fisher Scientific). After 48 h, viral supernatants were collected and centrifuged at 2000 rpm for 10 min to get rid of cell debris. The supernatant was filtered through a 0.45μm ultra-low protein binding filter (Merck Millipore). Aliquots were stored at −80 °C.

### Generation of ACE2 and CRISPR-activation cell line

HEK293 cells expressing human ACE2 (previously described^20^) were transduced with lentiviruses carrying dCAS9-10xGCN4 (Addgene 60903) termed HEK293-ACE2-SunTag. Cells were then sorted and seeded as single clones on 96-well plates. Clones were expanded and screened for their ability to drive expression of a positive target gene by CRISPR-activation. Expression was determined by qRT-PCR (**Supplementary Figure 1a**).

### Whole genome CRISPR-activation screen

HEK293-ACE2-SunTag cells were infected with the CRISPRa-V2 library at a multiplicity of infection (MOI) of 0.3, (3×10^7 cells, i.e, a minimum of 300 cells per guide) in T175 formats. Cells were selected with puromycin 1.6ug/ml for 3 days. CRISPR-targeted cells were then inoculated with authentic SARS-CoV-2 (Wuhan) at a dose leading to 90% cell death after 72 hours. Surviving cells were kept for an additional 72 hours with daily media changes. Additional control flasks were maintained throughout the same period without virus treatment. Cells were harvested for genomic DNA extraction. Screen was performed with 2 biological replicates per condition.

### Next generation sequencing

Genomic DNA corresponding to selected and diversity control samples were PCR amplified with NEBNext High-Fidelity 2X PCR Master Mix (New England Biolabs) as described in^20^. The following primers were used in this reaction: NGSCRISPRiaF1 5’-CAGCACAAAAGGAAACTCACCCTAACTG-3’ and NGSCRISPRv2Rev1 5’-TGTGGGCGATGTGCGCTCTG-3’. A second PCR reaction with staggered primer mix P5 and P7 indexing primer unique to each sample was prepared as previously described^40^. Reactions were isolated by gel electrophoresis and sent to the Ramaciotti Centre for Genomics for next-generation sequencing. Raw sequencing reads were mapped and counted using MAGeCK (v0.5.9.2)^41^. Read counts were then analysed using CRISPHieRmix^42^ to identify enriched and depleted genes as compared to diversity controls. See **Supplementary Extended Table 1** with gain-of-function and loss-of-function table for a full list of genes identified with CRISPhieRmix. The top 100 ranked genes, as determined by Local FDR, were selected and uploaded to Ingenuity (v01-21-03) for core analysis (IPA). KEGG pathway analysis was similarly performed on the top 100 ranked genes.

### Cell line generation for target validation using authentic SARS-CoV-2 virus

SgRNA sequences for target genes were cloned into pLentiguide puro (Addgene 52963) and confirmed by sanger sequencing. A full list of target sequences is found in **Supplementary Table 2**. Viruses were generated as described above. HEK-ACE2-SunTag clone used in the activation screen was infected with LentiRNA and selected with Puromycin for a period of 3 days before screening guides for target activation. RNA was extracted following manufacturer’s protocol (Total RNA Kit Farvogen). **Supplementary Table 3** lists all sgRNA sequences used for CRISPR-activation. Cells were seeded as described in “Authentic SARS-CoV-2 virus in vitro assays” section below.

### Authentic SARS-CoV-2 virus in vitro assays

Suntag CRISPR-activation cells were plated onto 384 well plates. 24 hours later cells were inoculated with serially diluted SARS-CoV-2 isolates in culture medium, where an equal volume was added to each well. For transient expression, tests were carried in HEK-ACE2-TMPRSS2 cells generated as previously described^7^. These cells were transfected as described in “cDNA Transfection” and seeded 24-hours post-transfection. Cells were then inoculated as described with isolates of SARS-CoV-2 variants (D614G or Delta). Plates were incubated for 48 hours at 37°C before addition of NucBlue™ live nuclear dye (Invitrogen, USA) at a final concentration of 2.5%. After a 4-hour incubation plates were imaged using an IN Cell Analyzer HS2500 high-content microscopy system (Cytiva). Quantification of nuclei was performed with automated IN Carta Image Analysis Software and normalised to uninfected wells. Genes that did not reach statistical significance in the validation are reported in **(Supplementary Fig. 1d)**.

### Cell line generation for spike binding screening

SgRNA sequences for target genes were cloned into pXPR502 (Addgene 96923) and screened by sanger sequencing. A full list of target sequences is found in table 1. Viruses were generated as described above, and transduced in a HEK cell clone expressing CRISPR-activation (SAM) (addgene 113341) previously described in^20^. Cells were incubated for a period of 3 days with 2ug/ml of puromycin before testing screening guides for SARS-CoV-2 spike binding capacity as described in “Flow Cytometry - Spike binding protein assays” section.

### SARS-CoV-2 Spike labelling and conjugation

The expression construct used to generate SARS-CoV-2 spike protein was a gift from Jason McLellan (Addgene #154754). Spike protein was prepared as previously described in^20^. Spike protein was conjugated to Alexa Fluor™ 647 using a protein labelling kit (Cat A20186, Invitrogen) according to manufacturer’s instructions. Briefly, a solution of 50 μL of 1 M sodium and 500 μL of 2mg/mL protein was prepared and incubated at room temperature for an hour. Conjugated protein was loaded onto Bio-Rad BioGel P-30 Fine size exclusion purification resin column and eluted by centrifugation. Protein quantification was determined using NanoDrop (ThermoFisher Scientific).

### Flow Cytometry - Spike binding protein assays

Spike binding assays were performed as described in^20^. Cells were dissociated with TryplE for 5 mins at 37C and then washed and then washed with culture media by centrifugation. A minimum of 100k cells were incubated with Alexa Fluor 647-conjugated SARS-CoV-2 spike glycoprotein (50ug/ml) for 30 mins at 4°C. Cells were then washed with 1% BSA in PBS, and resuspended in the same solution before analysis with the Cytek Aurora (Cytek Biosciences). 3 independent experiments were performed for these assays.

### Pseudotyped virus generation and DiD-labelling

Pseudotyped lentiviruses were generated as previously described^20^. Briefly, pseudoviruses were generated using a five-component plasmid system. SARS-CoV-2 (Wuhan) and Delta were made as described^20^. Pseudotyped MERS and SARS-CoV-1 receptor protein sequences were cloned onto the same backbones as of SARS-CoV-2 using NEBuilder HiFi DNA Assembly Master. DiD labelling was performed by adapting from a previous protocol^43^. Media containing virus particles was first concentrated to 20x using 100K MWCO protein concentrator columns (Thermofisher). Concentrated virus was then incubated using 1:2000 DiD-Cell labelling solution (ThermoFisher V22887) for 20 mins at 37°C. Medium was then diluted by a factor of 10, and reconcentrated using Polyethylene Glycol <u>8000</u> (PEG) overnight at 4°C. The next day the media was centrifuged at 3200G for 30 mins at 4°C. The supernatant was discarded and the pellet containing labelled viral particles was resuspended in DMEM and aliquoted. Virus was titered using the QuickTiter™ Lentivirus Titer Kit (Cell Biolabs, Inc) following manufacturer conditions.

### DiD labelling of authentic SARS-CoV-2

SARS-CoV-2 (Wuhan) isolates were expanded in VeroE6-TMPRSS2 cells as described^44^. For labelling, medium containing virus isolates was incubated in Dulbecco’s Modified Eagle Medium (DMEM; Gibco, 11995073) with 1:10000 Vybrant (V22887, Invitrogen) for 30 minutes at 37°C. This medium was resuspended with 1X PEG (ab102538, Abcam) and left overnight at 4°C. Medium was then centrifuged for 30 mins at 3200g at 40°C. The supernatant was removed and the virus was resuspended, aliquoted and stored at -80C until use.

### Platelet isolation and activation

Blood was obtained from healthy human donors in accordance with the Human Research Ethics Committee of the University of Sydney (2014/244) and the declaration of Helsinki. Platelet isolation was performed as described^45,46^. Briefly, whole blood was centrifuged at 200 G for 20 min. The supernatant (platelet-rich plasma) is collected and anti-coagulated further with citrate-dextrose. After incubating for 30 min at 37°C, the platelet-rich plasma is treated with 2μM Prostaglandin E1 (Sigma) and centrifuged at 800 G for 10 min. The supernatant was discarded and platelet pellet was resuspended in Hepes-Tyrode’s buffer. Platelets were tested within 2 hours of isolation. Platelets were activated using Thrombin (0.1 U/ml) (Sigma) for 7 mins at room temperature, and fucoidan (100nM) where relevant. CD62p-PE antibody (12-0626-82 Thermofisher), Spike-Alexa647 (10μg/ml), or DiD-Wuhan (3×10^^7^ particles/ml) was then added for an additional 7 mins. PPACK (60nM) was then added to stop Thrombin action. Platelets were then fixed in 4% PFA for 10 mins before proceeding for further analysis using flow cytometry. Platelet assays were performed from a minimum of 3 independent donors.

### HUVEC gene editing

Human umbilical vein endothelial cells (HUVECs) were sourced from consenting donors under Ethics approval by the Sydney Local Health District Human Ethics Committee, X16-0225. HUVECs were obtained from umbilical cords by collagenase treatment (32013974). Cells were maintained in MesoEndo growth medium (Lonza, Basel, Switzerland) in 5% CO2 incubator. HUVECs were cultured to 90% confluency. SELP-KO populations were generated by transduction with CRISPR-V2 lentivirus^5^ prepared as described above. The following sgRNA sequences were used for generating KO1, and KO2 populations (5’-GTCACAGATGAATTGACATG-3’, 5’- ATAGTTCGGTGTGATAACTT-3’, respectively). An intergenic control guide (5’- TGGCAACATATATAAGCAAG 3’) was used for control. Transduction was performed by an hour of centrifugation at 930G 31C in the presence of polybrene (8ug/ml). After spinfection the media was removed. 36 hours post-transduction cells were treated with 0.5ug/ml puromycin for 48 hours. Cells were then allowed to expand before subsequent assays. HUVEC characterizations were performed on a minimum of 3 independent donor cell lines.

### RNA extraction and RT-qPCR

RNA was isolated from cells using the ISOLATE II RNA Mini Kit (Bioline) and concentration was measured by Nanodrop (Thermo Scientific). cDNA was synthesised using the iScript Select cDNA Synthesis Kit (Bio-Rad) according to manufacturer’s instructions. Briefly, 50-500 ng of RNA was added to iScript RT Supermix and nuclease-free water to a final volume of 10 μL. The assembled reactions were then incubated in a thermocycler as follows: 25°C for 5 min, 46°C for 20 min and then 95°C for 1 min. RT-qPCR was then performed on the cDNA samples using SYBR Select Master Mix (ThermoFisher Scientific) and the LightCycler 480 System (Roche). All primer sequences used are listed in **Supplementary Table 4**. Results were analysed using the ∆∆CT method and are presented in (**Supplementary Fig. 1e**).

### cDNA transfection

A SELP-TurboGFP fusion construct was purchased from Origene (RG209822). TurboGFP control was generated by digesting SELP-GFP plasmid with MluI and BamHI, and ligating with annealed oligos (5’-CGCGCGAATTCATGCTAGCAACCGGTGCA-3’,

5’-GATCTGCACCGGTTGCTAGCATGAATTCG-3’). The ACE2-GFP fusion construct was generated using NEBuilder HiFi DNA Assembly. Briefly this construct was made by PCR amplification of the SELP-GFP vector (5’- ACGCGTACGCGGCCG-3’, 5’- GGCGATCGCGGCGGC-3’), and an ACE2-cDNA insert (5’- TGCCGCCGCGATCGCCATGTCAAGCTCTTCCTGGCTCC -3’, 5’-GCGGCCGCGTACGCGTAAAGGAGGTCTGAACATCATCAGTGT – 3’). Cells were plated 24 hours before transfection on 6 well plates (720k per well). Cells were transfected with respective constructs (2.4 μg/well) using Lipofectamine 3000 according to manufacturer’s protocols. Cells were replated for subsequent experiments after 48 hours of transfection.

### Microfluidics Assay

Microfluidic devices^45^ were coated overnight at 4C with fibronectin (100 ug/ml). The next day 20,000 HEK cells transiently expressing GFP, ACE2-GFP, or SELP-GFP loaded onto the inlet chamber in a volume of 10μl and were allowed to settle for 3 hours. Perfusion with DID-labelled pseudoviral particles at a shear rate of 100 s^-1^ for a period of 1 hour per chip was performed. All studies were completed within 4 hours after cells were settled. Chambers were washed with 100 ul of PBS, followed by 4% PFA fixation at room temperature for 20 mins. Chambers were then washed with 100 ul of PBS and stained with Hoescht for 10 mins, before a final wash with PBS. Slides were then imaged and quantified using Opera Phenix Plus (Perkin Elmer). Microfluidic assays were repeated as 3 independent experiments.

### Immunocytochemistry

Cells were fixed with 4% PFA for 20 minutes at room temperature. Cells were then blocked for an hour (PBS with 5% Normal Goat Serum, 1% BSA, 0.05% Triton-X, 0.3M Glycine). Cells were then incubated with respective antibodies (**Supplementary table 3**) in blocking solution for 1 hour at room temperature, followed by washes with PBS-0.05% Triton-X. Corresponding secondary conjugated antibodies and hoechst 33142 suspended in blocking solution was then used to incubate cells for an hour at room temperature. Cells were then washed with PBS-0.05% Triton-X, and PBS was added for imaging or placed on coverslip.

### High-throughput imaging

Cells were imaged using a Perkin Elmer’s Opera Phenix. High-throughput image analysis was performed using Harmony Software (Perkin Elmer).

### Flow Cytometry of labelled authentic SARS-CoV-2

Vero-E6-hACE2 and Hela cells were lifted with TryplE for 5 mins at 37°C and neutralised with culture media by centrifugation. 100k cells were resuspended in 20 μl of cell media containing DiD-SARS-CoV-2 at 0.5 and 1 MOIs or control media. Cells were incubated for 45 mins at 37 °C. Cells were then fixed with 4% PFA for 30 mins at room temperature, and washed twice with FACS buffer consisting of PBS, 2% BSA, 0.5mM EDTA. Cells were then analysed by flow cytometry using BD Canto.

### Mice inoculation

All experiments involving animals were approved by the University of Calgary Animal Care Committee (Protocol # MO8131) and conform to the guidelines established by the Canadian Council for Animal Care. Transgenic mice expressing human ACE2 (hACE2) were purchased from Taconic (B6;C3-Tg(CAG-ACE2)70Ctkt), and then bred in house. All mice were housed under specific pathogen-free, double-barrier unit at the University of Calgary. Mice were fed autoclaved rodent feed and water ad libitum. 8-12-week-old hACE2 mice were administered 2.5×10^4^ p.f.u. SARS-CoV-2 via intranasal route. Briefly, mice were anesthetised under 3% isoflurane (with oxygen as carrier), for 2-3 minutes, and 25 ul of SARS-CoV-2 diluted in PBS was inoculated onto the nostril, until the mouse inhaled the drop. After 4 days, mice were anesthetised (10 mg/Kg xylazine hydrochloride and 200 mg/Kg ketamine hydrochloride) and the jugular was cannulated for the administration of labelled SARS-CoV-2 (4.8 × 10^6^ p.f.u.). 3 mice were utilised per condition.

### Spinning Disk Confocal Intravital Microscopy

Stabilised pulmonary intravital microscopy was performed as previously described^47^. Briefly, anaesthetised mice (ketamine and xylazine) received a right internal jugular intravenous catheter to administer fluorescent antibodies, DiD-SARS-CoV-2, and/or Anti-P-selectin treatment (100 ug; rat anti-mouse CD62P, BD Pharmigen). To visualise the endothelium and platelets, 5 μg of fluorescently conjugated anti-CD31 (clone 390, BioLegend; either Alexa 594) and anti-CD49b antibodies (clone HMa2, BioLegend; Alexa 488), respectively, were administered intravenously 10 min before imaging. For traffic visualisation of virus, DiD-SARS-CoV2 (4.8 × 10^6^ p.f.u.) was administered. The trachea of the mouse was exposed for the insertion of a small catheter, which was then attached to a ventilator (Harvard Apparatus). The mouse was placed on its right lateral decubitus position and a small surgical incision was made. The intercostal muscles between ribs 4 and 5 was gently teased apart for the insertion of a lung window. The lung was then stabilised with a suction of 20 mmHg. Images were acquired with an upright microscope (BX51; Olympus) using a 20X/0.75 NA XLUM Plan F1 objectives (Olympus). The microscope was equipped with a confocal light path (WaveFx, Quorum) based on a modified Yokogawa CSU-10 head (Yokogawa Electric Corporation). A 512 × 512–pixel back-thinned EMCCD (electron-multiplying charge-coupled device) camera (C9100-13, Hamamatsu) was used for fluorescence detection. At least 3 fields of view with identified foci corresponding to DiD labelled SARS-CoV-2 were captured per animal. Images were acquired every ∼13s, for 10-30 minutes. Images were processed and analysed in ImageJ2 V2.9.0/1.53t.

### Intravital quantification of DiD-SARS-CoV-2

Video images were processed and analysed in ImageJ2 V2.9.0/1.53t (FIJI)^48^. Quantification was performed for the channel of interest by identifying foci using elliptical tools. Respective regions were then measured for grayscale intensity throughout the recorded period and exported as a table. Rate of change (rolling average) in intensity was measured by calculating the difference in intensity across 3 frames. Statistics were performed with GraphPad Prism 9. For individual foci tracking, videos were blinded and manually tracked using the Manual Tracking plugin, individual particles were then centred and plotted using R-Studio. Individual foci were also blindly evaluated to determine the number of new viral particles that were anchored throughout recordings.

### Quantification and statistical analyses

Statistical analysis was performed using GraphPad Prism 9 unless otherwise specified. All error bars in this manuscript report SEM. Imaging analysis and quantification for microfluidic studies and lentiviral work was performed using Harmony High-Content Imaging and Analysis Software (Perkin Elmer). All flow cytometry data was analysed using FloJo Software v10.6 (BD Life Sciences).

## Supplementary Information

**Supplementary Fig. 1.**
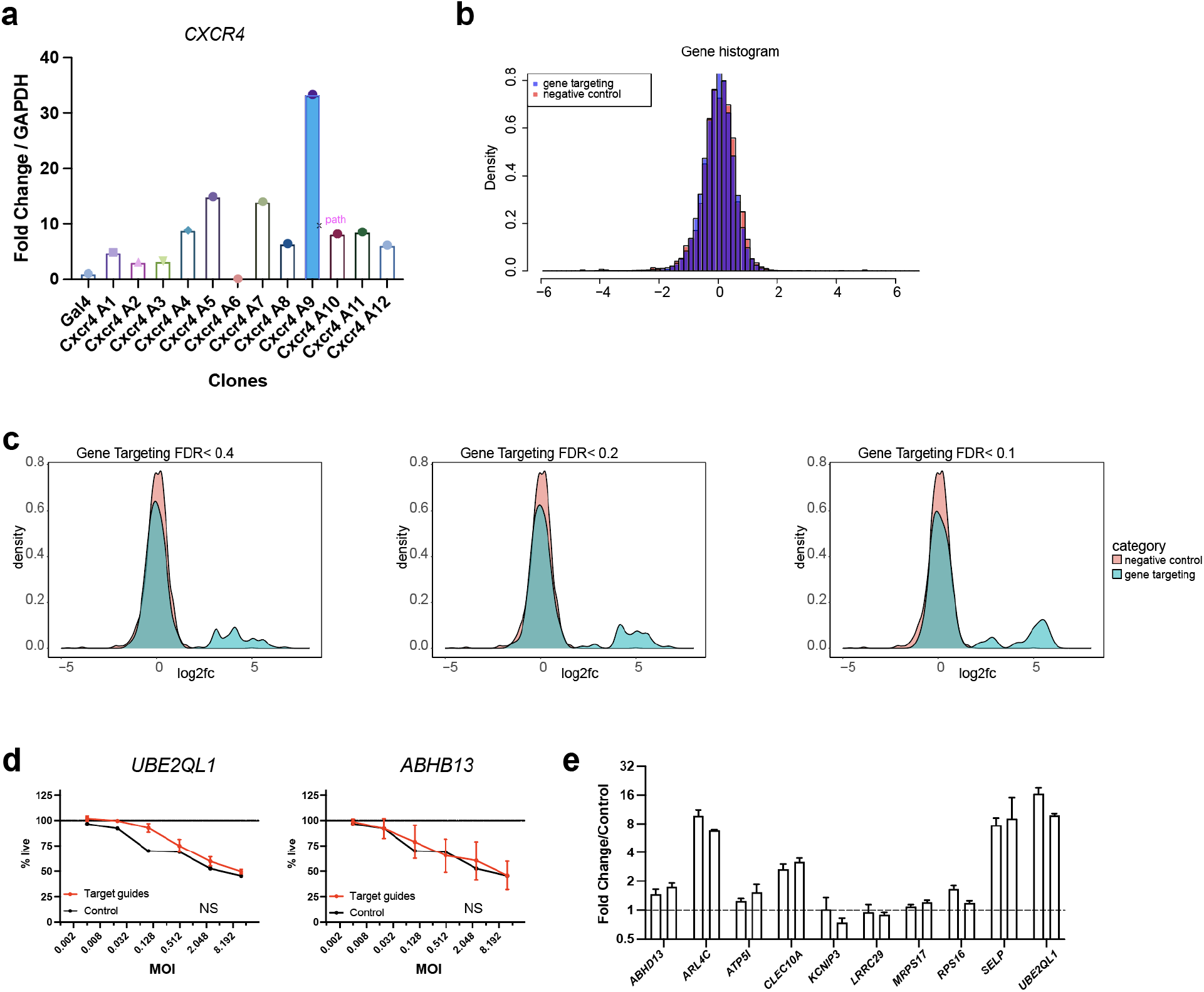
**a**, qRT-PCR screen showing expression of *CXCR4* by dCAS9-Suntag machinery in HEK-ACE2 cells. Blue bar indicates clone selected for future studies. *CXCR4* was chosen as a target gene based on previously reported sgRNA guides^1^. **b**, Superimposed histograms displaying negative control guides with targeting genes showing normal distributions. **c**, Density diagrams showing superimposed negative distribution vs. gene targets with decreasing False Discovery Rates (FDR). **d**, Individual target validation that did not show significant protection against SARS-CoV-2. **e**, Expression of target genes by each individual sgRNA and normalized to control negative guides.

**Supplementary Fig. 2.**
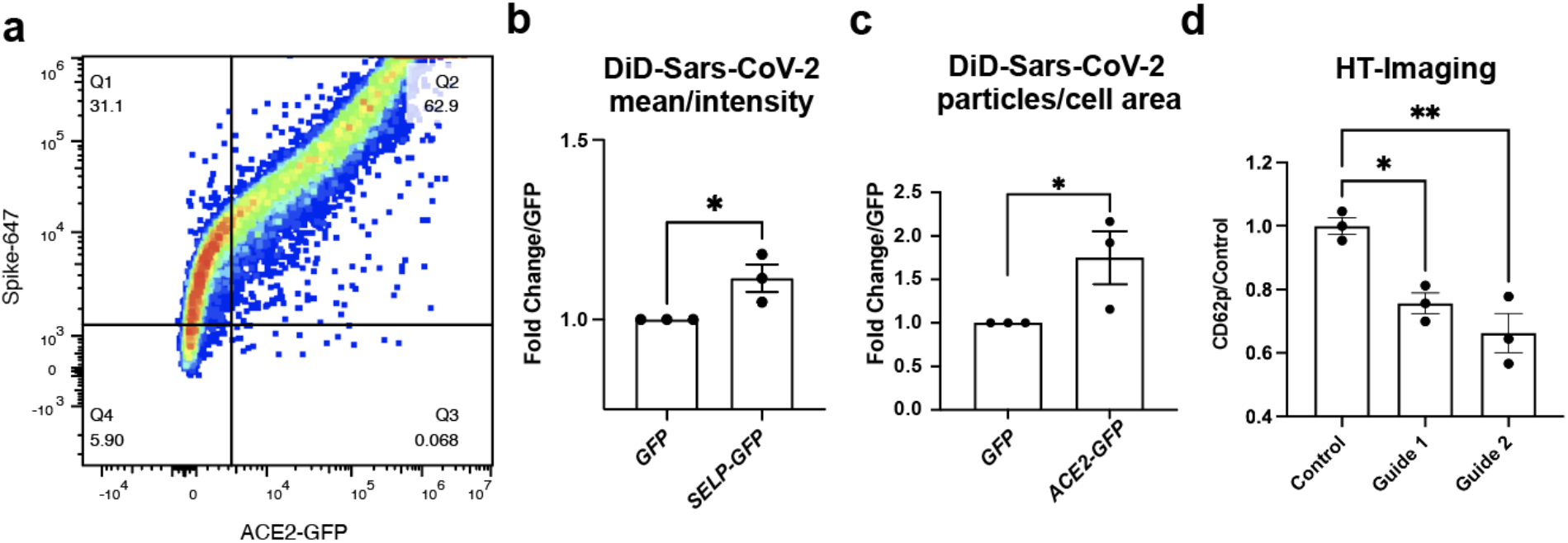
**a**, Sample flow-cytometry plot showing ACE2-GFP expressing cells binding SARS-CoV-2 Spike-647. **b**, Quantification of DiD-pseudovirus fluorescent intensity in microfluidic capillaries lined with SELP-GFP cells or controls. **c**, Quantification of DiD-pseudovirus fluorescent particles in microfluidic capillaries lines with ACE2-GFP or controls. Significance was determined by t test *,P<0.05. **d**, High-throughput imaging quantification of CD62p positive HUVECs after CRISPR knock out of SELP. Significance was determined by 1-Way ANOVA and dunnetts, **, P<0.01; *, P<0.05.

**Supplementary Fig. 3.**
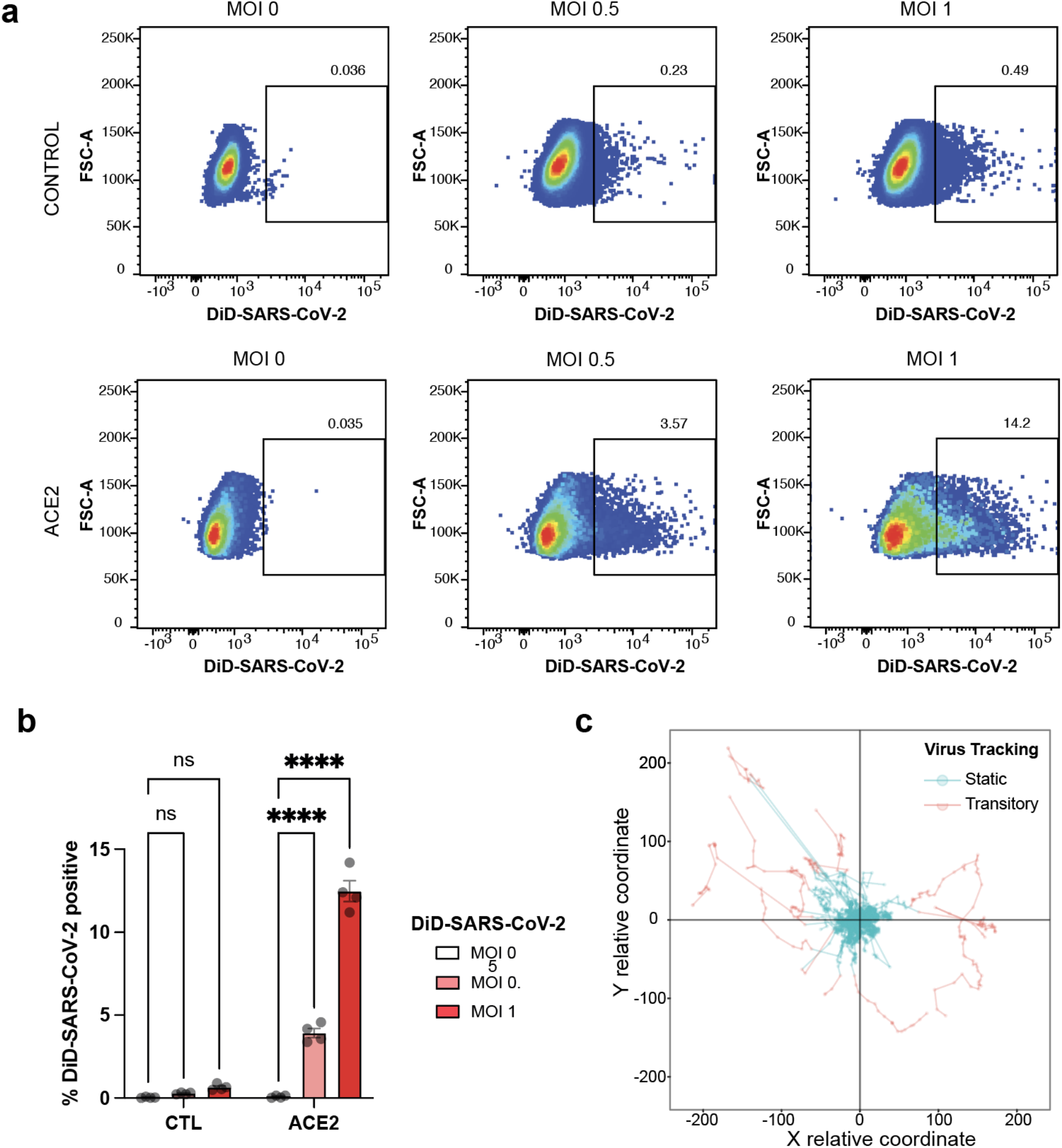
**a**, Sample flow-cytometry plot showing preferential binding of DiD-SARS-CoV-2 in control cells or Vero6-hACE2 cells. **b**, Percent of DiD-SARS-CoV-2 cells. Significance was determined by 2-Way ANOVA and Dunnets ****, P<.0001. **c**, Individual tracks of DiD-labelled particles, compared from a central origin point.

**Supplementary Table 1a.**
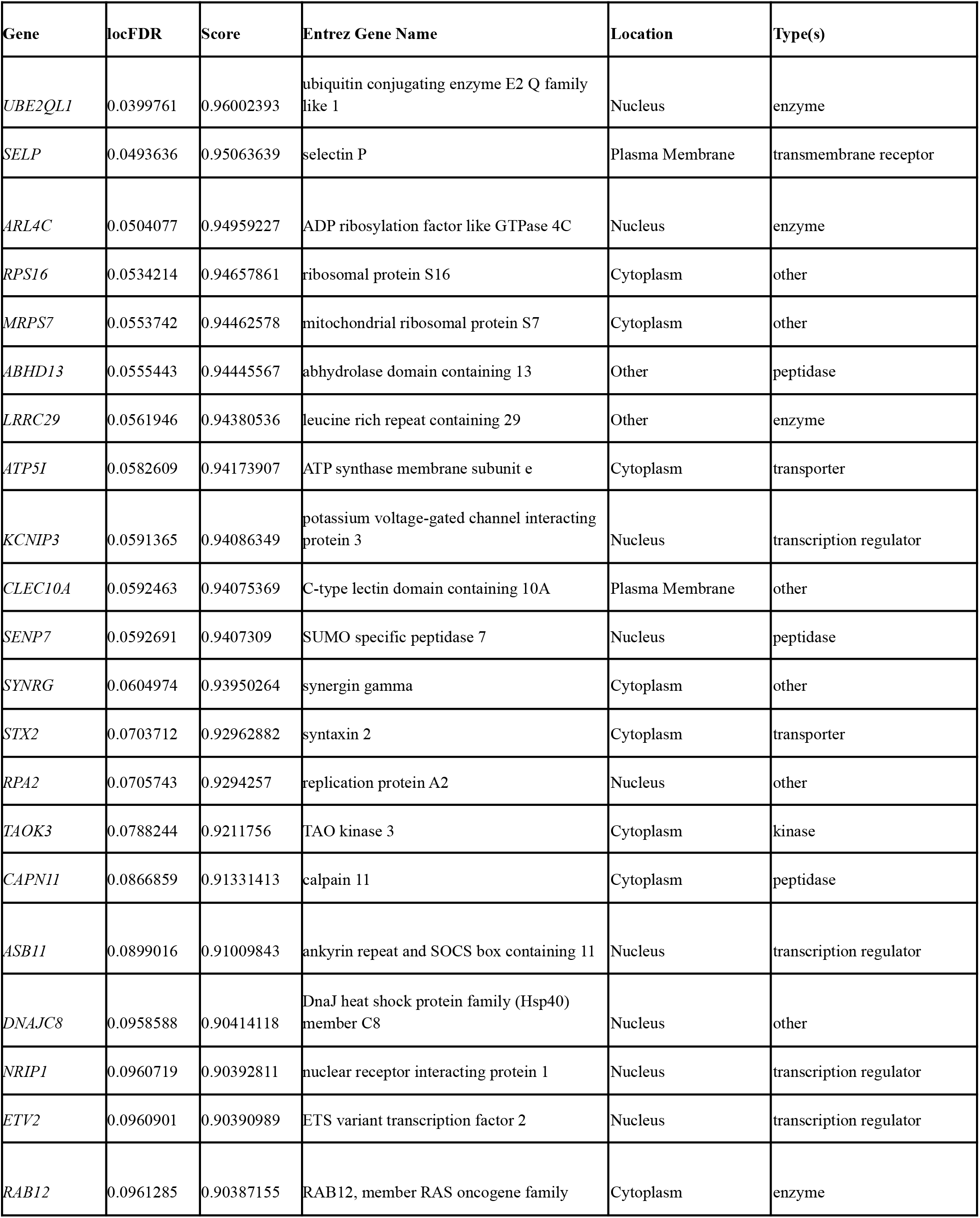
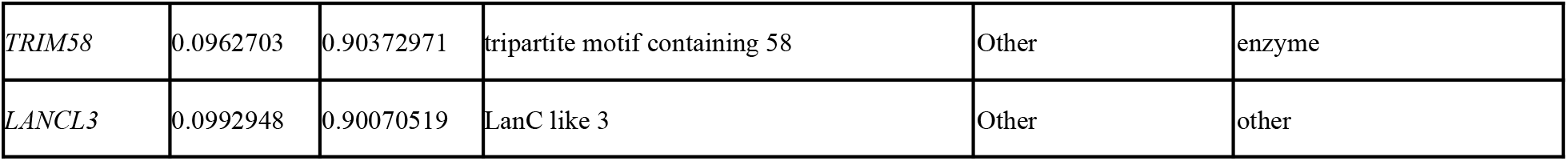
CRISPR-Activation screen for overrepresented genes with FDR <0.1. See extended table for a list of all genes.

**Supplementary Table 1b.** CRISPR-Activation screen for top 10 underrepresented genes with FDR <0.1. See extended table for a full list of genes.

| Gene | locFDR | Score | FDR |
| --- | --- | --- | --- |
| *ATP6AP1L* | 0.30500658 | 0.69499342 | 0.30500658 |
| *GFM2* | 0.48960427 | 0.51039573 | 0.39730543 |
| *PEPD* | 0.69577662 | 0.30422338 | 0.49679582 |
| *PICK1* | 0.69745181 | 0.30254819 | 0.54695982 |
| *COX7C* | 0.71498018 | 0.28501983 | 0.58056389 |
| *GPKOW* | 0.71777868 | 0.28222132 | 0.60343302 |
| *MMP3* | 0.72229919 | 0.27770082 | 0.6204139 |
| *KLK1* | 0.72337579 | 0.27662422 | 0.63328414 |
| *UTP18* | 0.7317411 | 0.2682589 | 0.6442238 |

**Supplementary Table 3.**
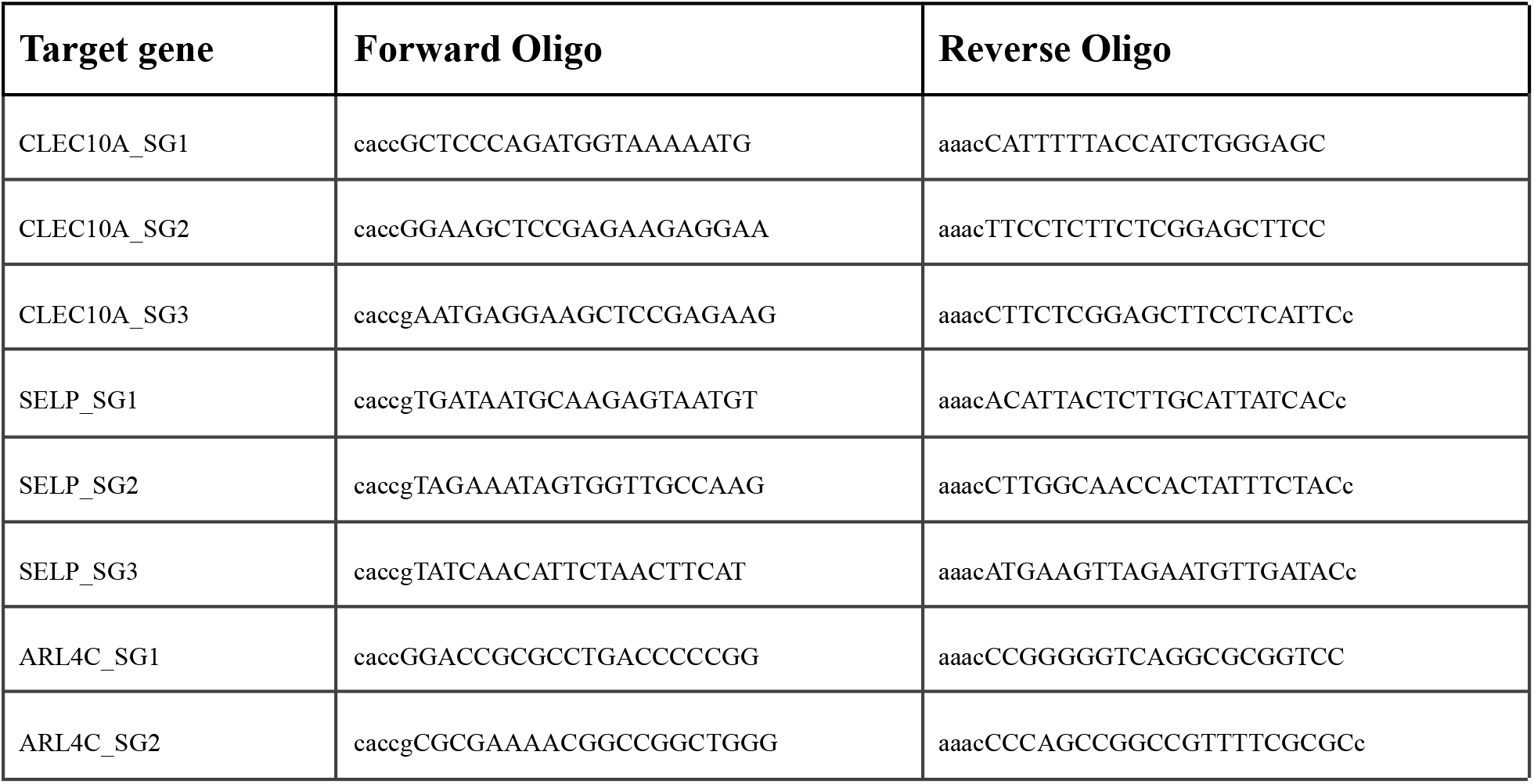
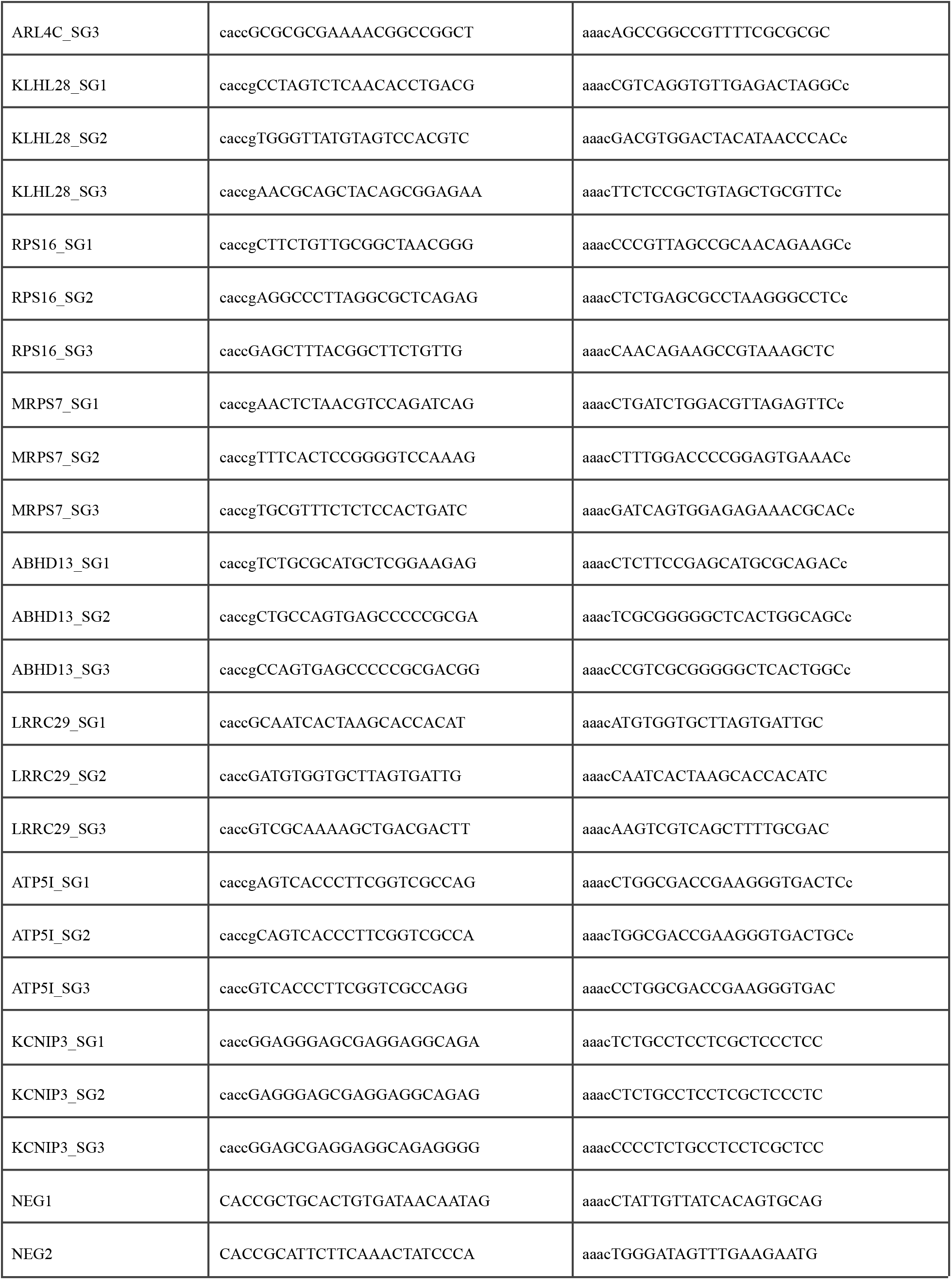
CRISPR-Activation sgRNA oligos cloned for individual target evaluation.

**Supplementary Table 4.**
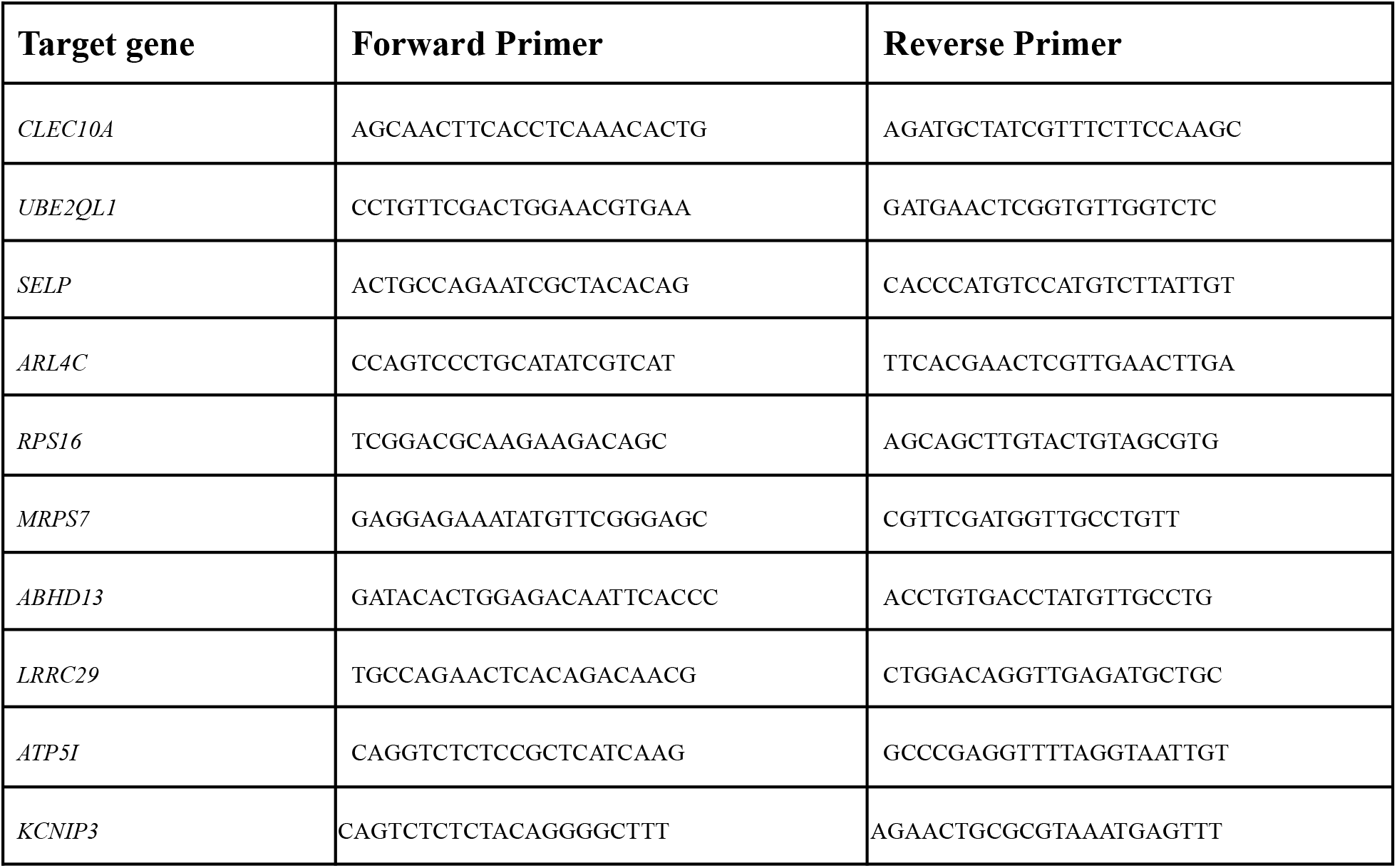
Primers used in RT-qPCR.

**Table 3.**
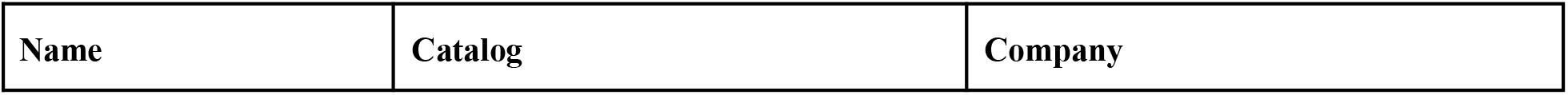
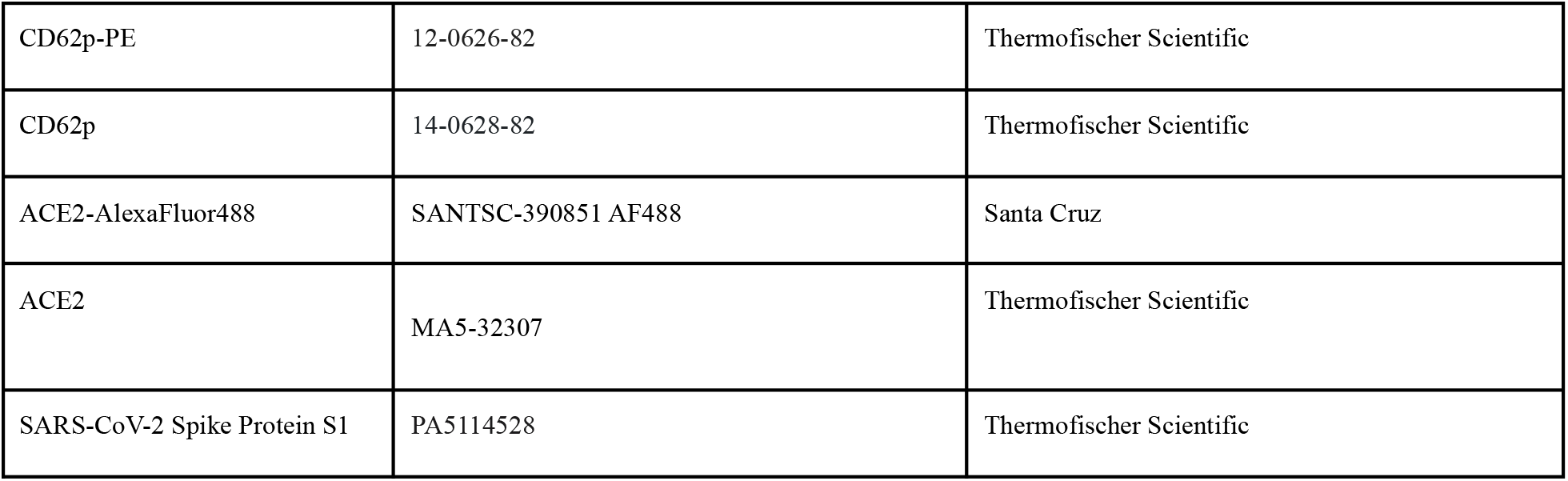
Antibodies

| **Name** | **Catalog** | **Company** |
| --- | --- | --- |
| CD62p-PE | 12-0626-82 | Thermofischer Scientific |
| CD62p | 14-0628-82 | Thermofischer Scientific |
| ACE2-AlexaFluor488 | SANTSC-390851 AF488 | Santa Cruz |
| ACE2 | MA5-32307 | Thermofischer Scientific |
| SARS-CoV-2 Spike Protein S1 | PA5114528 | Thermofischer Scientific |

## Notes

### Competing Interest Statement

The authors have declared no competing interest.

